# Closing Gaps But Increasing Bias In North American Butterfly Inventory Completeness

**DOI:** 10.1101/2020.07.20.212381

**Authors:** Vaughn Shirey, Michael W. Belitz, Vijay Barve, Robert Guralnick

## Abstract

Aggregate biodiversity data from museum specimens and community observations have promise for macroscale ecological analyses. Despite this, many groups are under-sampled, and sampling is not homogeneous across space. Here we used butterflies, the best documented group of insects, to examine inventory completeness across North America. We separated digitally accessible butterfly records into those from natural history collections and burgeoning community science observations to determine if these data sources have differential spatio-taxonomic biases. When we combined all data, we found startling under-sampling in regions with the most dramatic trajectories of climate change and across biomes. We also found support for the hypothesis that community science observations are filling more gaps in sampling but are more biased towards areas with the highest human footprint. Finally, we found that both types of occurrences have familial-level taxonomic completeness biases, in contrast to the hypothesis of less taxonomic bias in natural history collections data. These results suggest that higher inventory completeness, driven by rapid growth of community science observations, is partially offset by higher spatio-taxonomic biases. We use the findings here to provide recommendations on how to alleviate some of these gaps in the context of prioritizing global change research.

## INTRODUCTION

The mobilization of openly and freely available natural history data has increased the ability for researchers to access information about species distribution and abundance in a given time and place. In recent years, these data have been augmented by community science programs which facilitate collection of biodiversity observations and digital vouchers from a network of volunteers. Aggregated data from both natural history collections and community science programs have been used to answer often broad questions in ecology, including assessing extinction risks for understudied groups (Carlson *et al*. 2017, Seppälä *et al*. 2018) and modeling species response to environmental change (Eskildsen *et al*. 2015).

Despite the utility of these data, many taxa are still under-sampled (Troudet *et al*. 2017) and prevalent biases in the spatiotemporal distribution of these data are noteworthy (Beck *et al*. 2013, Meyer et al. 2015). These biases imply that inventory completeness (how many species have been recorded vs. how many are expected to occur) is uneven across time and space. Given the urgency to understand ecological responses to many global change processes, knowing where sampling has and has not occurred to a sufficient degree is critical for both prioritizing effort to close information gaps and choosing extents and scales for macroecological analyses. The enormous growth of community science reporting for some groups promises to rapidly close inventory gaps, but less is known about how specimens from natural history collections and community science data may be differentially spatially biased. Community science volunteers may stay closer to developed areas to sample biodiversity than collectors who may be more attentive to collecting in under-sampled regions. This may lead to larger under-sampling by community scientists in remote regions, including the far North, which is projected to experience the most dramatic effects of climate change. Under-sampling in the Arctic and other sparsely populated regions negatively impacts ability to assess how climate has impacted communities over time.

Butterflies (Lepidoptera: Papilionoideae) are a diverse group of organisms that are relatively less sampled compared to vertebrate fauna (Troudet *et al*. 2017) which have been the focus on previous sampling completeness assessments (Meyer *et al*. 2015). Additionally, butterflies have been widely used to detect signals of global change (Parmesan *et al*. 1999, Eskildsen *et al*. 2015). Given the value of butterflies as an indicator group, we aim to test how well sampled North America is for butterflies using natural history collections and community science data, as gaps in openly accessible biodiversity data limit efforts to address ecological, evolutionary, and conservation questions. More specifically, we utilize estimates of distributions from field guides to establish a baseline richness value at multiple, coarse scales usable for presence prediction (Jetz et al. 2012). We then compare that value to richness derived from occurrence records from the Global Biodiversity Information Facility (GBIF), iDigBio, and eButterfly.

We separated occurrence records into those from natural history collections and from community science-based observations and examined temporal trends in the number of records and completeness for each. We then tested the hypothesis that both types of occurrences were biased to areas where the humans are likely to be most active, but that those biases were particularly severe for community science records. We also examine if there are differences across butterfly families among these record types, presuming that records from natural history collections are less likely to be biased in familial completeness coverage. To provide further context for these results, we ask how biomes and climate regimes are sampled differently in order to provide meaningful information for global change ecologists and other users of these data. Finally, we discuss potential strategies to mitigate under-sampling across the continent in the future.

## MATERIAL AND METHODS

Occurrence records were obtained from GBIF (GBIF 2020), iDigBio (iDigBio 2020), and eButterfly (Prudic *et al*. 2017) from 1950 through 2019 within North America (Canada, Mexico, United States). Range maps of species found in the United States and Canada were digitized from the *Kaufman Field Guide to Butterflies of North America* (Brock and Kaufman 2006). For species found in Mexico, range maps were digitized from *A Swift Guide to Butterflies of Mexico and Central America* (Glassberg 2018) as part of the ButterflyNet project, which are digitally available for visualization on Map of Life (Jetz *et al*. 2012). These maps only include known source population locations, and do not include distributions of strays. All range maps from the sources were merged into a single shapefile consisting of many spatial polygons which were clipped to only terrestrial areas within North America. These range maps were then intersected with continent-wide equal area grids at 100km, 200km, and 400km resolution. A species was considered to occupy a 100km cell if its range passed within 2,000m of the grid centroid and considered to occupy a cell at coarser resolutions if it intersected the grid cell irrespective of distance to the cell centroid. Taxonomic names across the fishnet grids and occurrence data were harmonized to a single taxonomic list using R package taxotools (Barve 2020) and the small minority of names that could not be resolved manually after the process were discarded from the analysis. We analyzed only occurrence records that fell within the boundaries of their species’ range map but recorded how many records fell outside of these boundaries over time to assess any potential temporal degradation of range maps.

Sampling completeness was calculated as the ratio of species observed in occurrence data within a grid cell to the number of overlapping range maps within that grid cell. In some cases due to range map exclusion along coastlines and because we only included species present in this fishnet if it occurred within 2000m of the grid centroid, this ratio was slightly higher than 1.0 and was thus floored to 1.0. The occurrence dataset was then filtered by the basis of the record, year, and taxonomy attributes to examine how specimen-only (listed as preserved specimen or material sample), community observation-only (listed as human observations from the basisOfRecord field in Darwin Core), time period, and the taxon-rank of family (which are monophyletic, Espeland et al., 2018) impacted completeness scores. Machine observations were a small fraction of these data and were not included in the analysis.

Overall average completion between specimen and observation data was assessed using a t-test. We then tested average completion differences among families using an ANOVA on the combined, specimen, and community observation datasets, and differences in the number of cells complete at or over 50% using a Chi-square test for families between specimen and observation based datasets. Post-hoc testing was conducted with Bonferroni correction in the case of Chi-square.

We also assembled spatial data including velocity of climate under RCP 8.5 forecasts into 2085 (AdaptWest 2015); human footprint, representing areas where there are built environments, roads, or converted land (Venter *et al*. 2016); protected regions (Dept. of Forestry and Natural Resources, Clemson University for CEC 2010); and biomes as designated by the World Wildlife Fund (Olson *et al*. 2001). For human footprint and climate velocity, we calculated average values, and for protected areas, the percent coverage of those areas, within each 100km grid cell. For biome type, we determined the majority biome within each 100km grid. We used these resampled values alongside the completion scores to identify drivers of sampling completeness and under-sampled regions described in more detail below.

For potential drivers of completeness, we considered human footprint and protected areas to each represent areas where humans may be actively reporting butterfly occurrences, and specified separate linear models for the combined, museum specimen, and community observation datasets as (Sampling Completeness ∼ Human Footprint + Protected Region Cell Coverage). We also ran these univariate models using either human footprint or protected areas as predictors. Model selection was then performed using AIC as the selection criterion to determine the top model. We compared model goodness of fit for the best models for natural history versus community science in order to assess the differential impact these factors may have on datasets with potentially different underlying observation strategies.

Finally, we examined the sampling completeness within the cells with the most extreme 10% and 25% of climate velocities and the sampling completeness across the WWF biomes found in North America. We removed from our analysis biomes in which the number of 100×100km cells was less than 10. This included mangrove forests, tropical grasslands, and flooded grasslands. All data preparation and analysis was performed in R version 3.6.3 “Holding the Windsock” (R Core Team 2020) using the packages tidyverse, sp, sf, raster, data.table, mapdata, maptools, gridExtra, stringr, rgdal, ggforce, exactextractr, and scales (Pebesma *et al*. 2005, Auguie 2017, Brownrigg 2018, Pebesma 2018, Dowle and Srinivasan 2019, Pedersen 2019, Wickham 2019a, Wickham 2019b, Baston 2020, Bivand 2020, Bivand and Lewin-Koh 2020, Hijmans 2020, Wickham and Seidel 2020). The script utilized here is available from a public GitHub repository at [anonymized]. It is also available with our generated datasets via a Zenodo archive at [anonymized].

## RESULTS

We obtained approximately 2.8 million records from our aggregate iDigBio, GBIF, and eButterfly datasets. Overall, 91.2% of occurrence records fell within range map delineations for their respective species. This has changed little over time with an average annual percentage of 88.6% from 1950-2019 and a recent increase within the last decade of sampling to 91.4%. From 1950 to 2019, the ratio of cells sampled biyearly at 80% completion by museum specimen data to those by community observations alone decreased dramatically, especially in the last decade of sampling with community based completion becoming more prevalent as the number of community observations increases (Figure 1).

**Figure 1.**
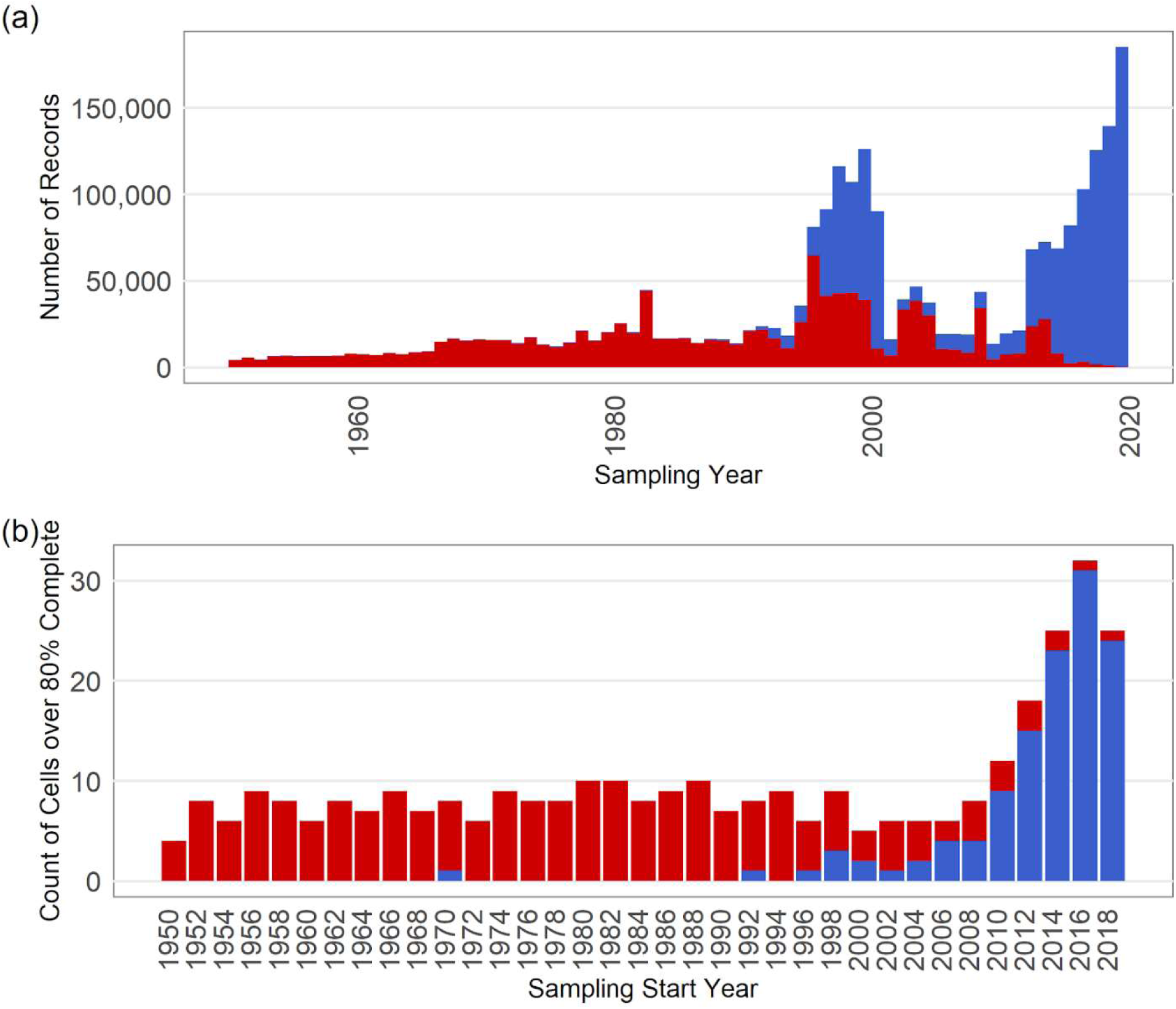
(a) The number of museum specimens and community observation-based occurrence records over time, stacked by year. (b) Number of cells at 100km resolution that are over 80% complete and that meet that threshold by museum or community observation data alone biyearly. Red = museum specimens; Blue = community observations.

### Human Footprint and Protected Areas

In all cases, the best performing model according to AIC included human footprint alone without the percentage of protected natural areas (Table 1). For museum records, the variance explained by the model was low (R^2^ = 0.09) compared to the composite dataset (R^2^=0.25) and the community science dataset (R^2^=0.29).

**Table 1.** Coefficient estimates of each multiple regression model for the full record set, specimen-only record set, and community observation recordset. Delta-AIC values indicate the difference between the multiple regression and simple regression model which included only human footprint as a predictor variable. In all cases, models excluding protected area percentage outperformed the simple regression according to AIC.

|  | *estimate* | *se* | *t* | *p-value* | *R<sup>2</sup>* | *delta-AIC* |
| --- | --- | --- | --- | --- | --- | --- |
| <u>All Records</u> |  |  |  |  | 0.25 | 1327.03 |
| **Intercept** | 0.354 | 0.0083 | 42.63 | < 0.0001 |  |  |
| **Human Footprint** | 0.027 | 0.0011 | 23.92 | < 0.0001 |  |  |
| <u>Specimens</u> |  |  |  |  | 0.09 | 1160.43 |
| **Intercept** | 0.194 | 0.0089 | 21.57 | < 0.0001 |  |  |
| **Human Footprint** | 0.014 | 0.0012 | 12.14 | < 0.0001 |  |  |
| <u>Observations</u> |  |  |  |  | 0.29 | 1144.99 |
| **Intercept** | 0.238 | 0.0088 | 26.96 | < 0.0001 |  |  |
| **Human Footprint** | 0.028 | 0.0011 | 24.79 | < 0.0001 |  |  |

### Geographic and Taxonomic Sampling Completeness

Sampling completeness was not homogeneous across scales with noticeable geographic gaps in the far north, midwest, and northern Mexico as illustrated in Figure 2. Mean specimen and observation-based completeness was significantly different according to our t-test (−13.27, 2919 DF, p < 0.0001), with observations having a higher average completion ratio (0.40 +/− 0.007 SE to 0.27 +/− 0.006 SE). Sampling was also inconsistent across families, especially within the Lycaenidae. To illustrate this better, in the composite dataset, differences among completeness across families were significant according to ANOVA (F_(4, 7267)_=51.49, p < 0.0001) (Figure 3a) and ANOVA also supported significant differences across families for the specimen based (F_(4, 5368)_=86.44, p < 0.0001)(Figure 3b) and observation based (F_(4, 6325)_=44.72, p < 0.0001)(Figure 3c) datasets (Post-hoc test results in Supplemental Figure 1). Chi-square tests to assess differences in the number of 100×100km cells completed at 50% or more between specimens and observations revealed there is a significant association with family-level completion and basis of record (X^2^=31.04, 4 DF, p < 0.0001)(Figure 3b,c). Post hoc comparisons (Beasley and Schumacker 1995) revealed that this association was significant for Nymphalidae and Pieridae with observations having more cells at 50% or more complete in these families (p < 0.01)(Figure 3d).

**Figure 2.**
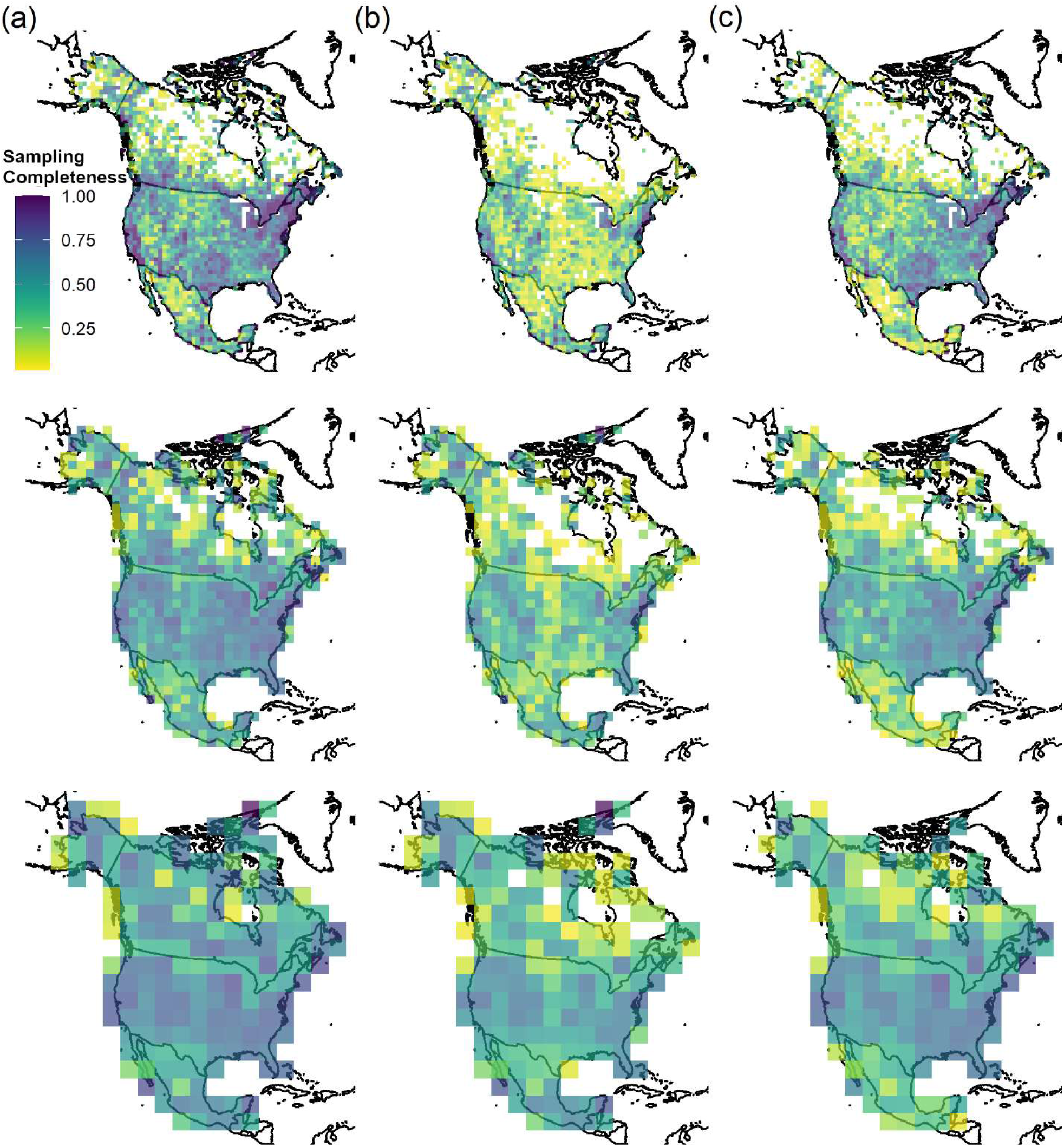
Sampling completeness within cells of varying spatial resolution (100km, 200km, 400km) across North America from 1950-2019 based on record source (a) all records, (b) specimens, and (c) community observations.

**Figure 3.**
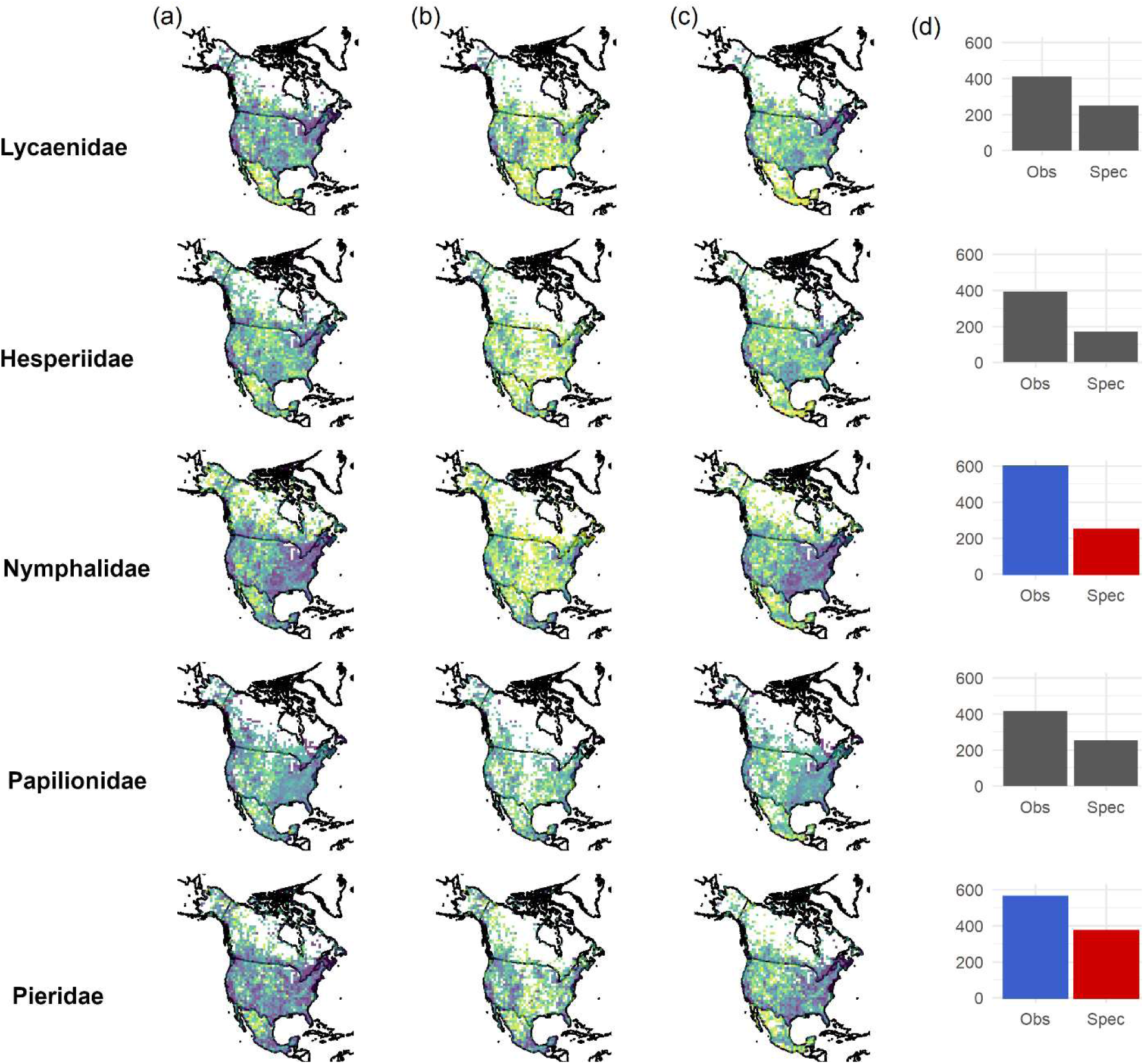
Sampling completeness among butterfly families at 100×100km resolution across North America from 1950-2019 by family and by record source (a) all records, (b) specimens, and (c) community observations. Panel (d) illustrates the number of cells over 50% complete in each family, colored plots indicate a significant contribution to the chi-square statistic after Bonferroni corrected post-hoc tests.

### Sampling in Projected Novel Climate Regimes and Biomes

Of the 80th and 95th percentile 100km resolution grid cells experiencing the most dramatic climate effects on average under RCP 8.5 into 2080, 97.5% and 97.2% fell below the 80% sampling completeness mark respectively, indicating under-sampling in these regions (Figure 4). In addition, sampling across biomes at the 100×100km resolution was inconsistent, with some biomes being sampled on average more completely than others as illustrated in Figure 5. Only the Mediterranean woodland/scrub biome demonstrated over 80% sampling completeness on average with notable under-sampling occurring in deserts, tropical, and boreal/arctic regions. Moderate sampling (between 50% – 80% completeness on average) was demonstrated within most mid-latitude temperate regions.

**Figure 4.**
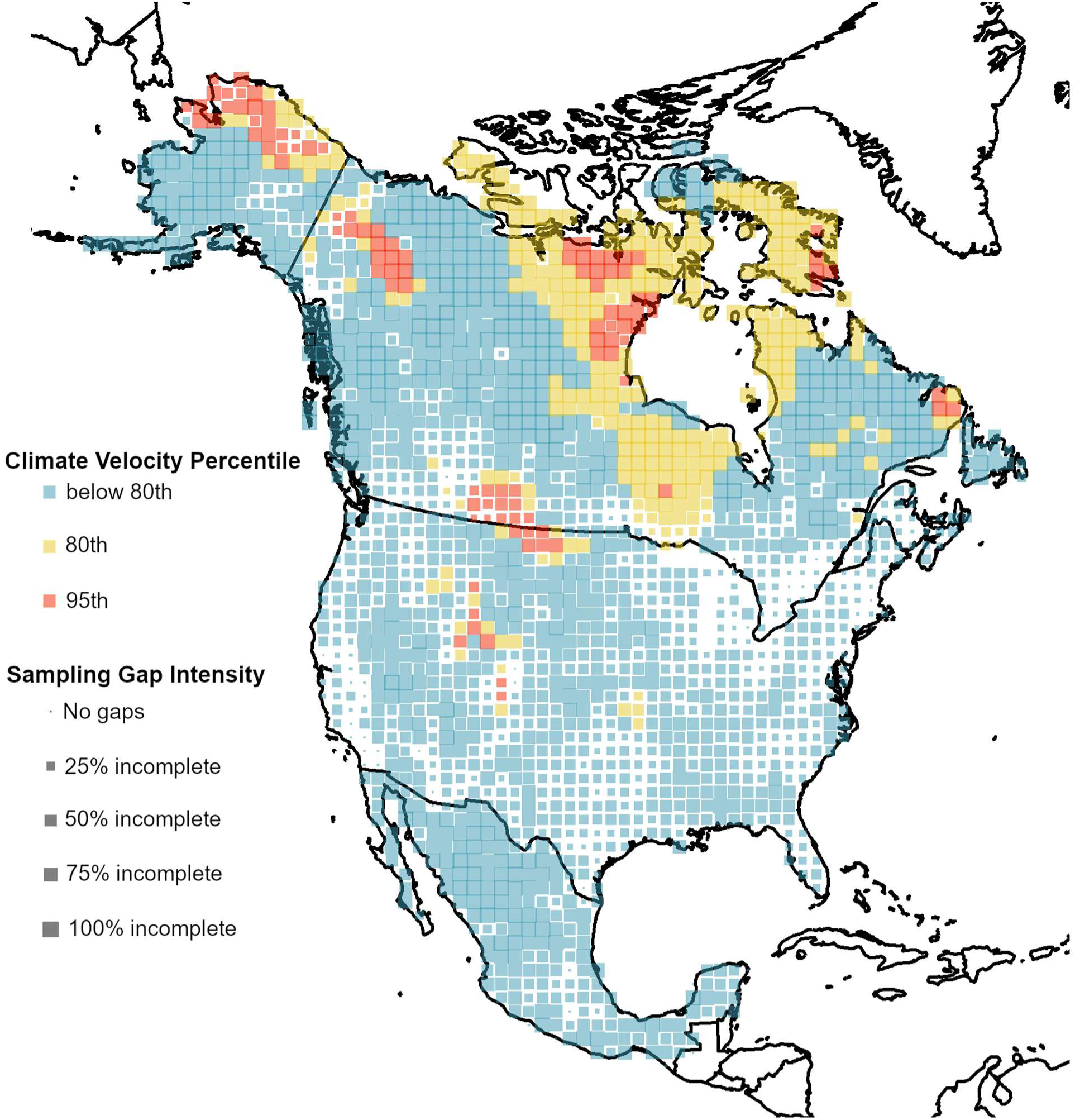
Sampling completeness of 100×100km grid cells by climate velocity percentile. Size of the point within the cell indicates sampling <u>incompleteness</u> (larger cells are less sampled). Yellow and red cells are the 80th and 95th percentile of climate velocities respectively. Blue cells fall underneath the 80th percentile for climate velocity. Climate velocity rasters do not extend into northern Nunavut.

**Figure 5.**
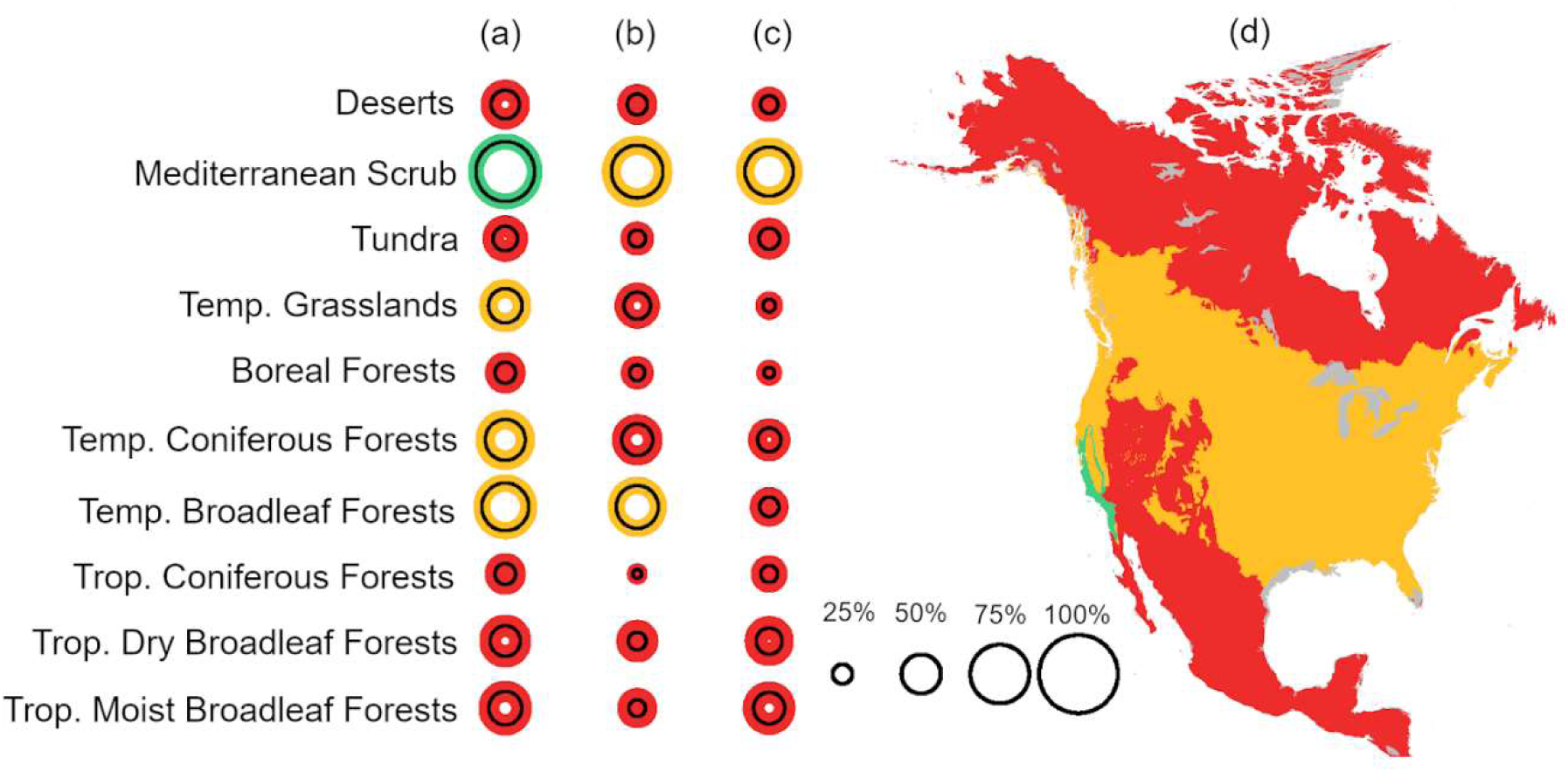
Mean +/− SD sampling completeness across WWF biomes, (a) composite dataset, (b) community observations only, and (c) museum specimens only. Panel (d) displays the biomes utilized without delineation for clarity and includes coloration based on average composite sampling completeness. Red = sampling below 50% average completeness, Yellow = sampling average between 50% and 80% completeness, Green = sampling average at or above 80% completeness. Grey regions represent unsampled areas, or regions where the number of 100×100km cells within a biome was < 15.

## DISCUSSION

Sample completeness across North American has accelerated in recent years, driven strongly by the growing number of community observations generated from programs such as iNaturalist (observations cited in our GBIF download), who share research grade observations with GBIF, and eButterfly (Figure 1a). The majority of cells with >80% completeness are now from community science data, which continues to grow exponentially per year (Figure 1b), demonstrating the importance of these data for closing distribution knowledge gaps into the future. A large volume of community science records may be due to the ease of submission. For example, iNaturalist submissions can be completed by simply taking a photograph on a mobile phone. Networks such as eButterfly often appeal more directly to dedicated amateur lepidopterists, and do not require photo vouchers to publish data, which has the potential to allow for more observations of butterfly species that are difficult to photograph. This is in contrast to specimen-based data in which preparation, curation, and digitization are all required steps to publish occurrence data.

Despite this influx of community science data, sampling is still inconsistent across space and taxonomy (Figure 2, 3). Regions with low human footprint are frequently under-sampled or not sampled at all, and our simple model validates this finding alongside other studies that have examined the relationship between human population densities and record densities (Girardello *et al*. 2019). A key finding is that these biases towards sampling where human infrastructure is the most developed are stronger for community observation data than for specimens (Table 1). Thus, community science observations are not likely to be a full panacea for closing inventory knowledge gaps. While some areas of North America are likely to be inventoried at increasingly finer spatial grain with burgeoning growth of community science data, other areas may remain perniciously under-sampled. This likely continuing butterfly inventory knowledge gap in remote regions is thus both particularly challenging and crucial to overcome since these are exactly the areas forecasted to experience the most climatic change. A particularly good example are polar regions of North America, where climate velocities are often particularly high (Figure 4 shows the 80th and 95th percentile of highest velocities) and sampling is woefully incomplete. (Figure 4). As well, even some mid and low-latitude biomes are under-sampled, including deserts and many tropical biomes in which butterfly diversity is extremely high (Willig *et al*. 2003) (Figure 3). We argue that community science alone is unlikely to solve existing gaps in biodiversity monitoring unless those programs are directed into sparsely populated regions through socially responsible excursions or other research campaigns that consult with local stakeholders and Indigenous communities. These directed and collaborative efforts, requiring partnerships and coordination, will help to provide a critical basis for mapping and ultimately monitoring butterfly diversity in order to detect changes in the face of shifting climate regimes.

We had anticipated that traits that make some butterflies easier to detect, photograph and identify might be biased across butterfly families, thus leading to familial-level biases in completeness. We expected these issues to be more acute for community scientists, compared to professional collectors, who presumably are collectively more knowledgeable and trained in sampling methods that might reduce bias. We already demonstrated reduced spatial biases for natural history specimen collecting, which might also suggest better sampling of habitats, also potentially reducing taxonomic biases. In the composite dataset, Lycaenidae exhibit lower average completeness with most other groups differing from each other as well (Supp. Figure 1), supporting our hypothesis of taxonomic biases in completeness. However, we did not find evidence that natural history specimen collecting led to less taxonomically biased sampling, at least at the familial level. We did however find that completeness from community science observations was higher compared to natural history specimen records only for nymphalids and pierids and not for other butterfly families. While higher completeness itself is not surprising given the trends we report, we had anticipated either similar trends across groups, or that showy groups such as the swallowtail butterflies (i.e. Papilionidae) were more likely to be biased in favor of community science observations given they are generally colorful, large, and charismatic. Further exploration using species-level trait data to tease apart these patterns is warranted. In particular, it may be that species-level rarity may be particularly important, especially if phylogenetically conserved. Other traits that may be worth examining include habitat and flight preferences (canopy vs. understory fliers) that directly relate to ease of human observation.

Our study expands upon prior work done on butterfly inventory completeness (Girardello *et al*. 2019) by including an independent baseline richness via digitized maps at coarse resolution and by examining the contributions of specimens and community observations. In addition, with a narrower focus on just North America and by including an assessment of sampling completeness in regions with high climate velocity and across biomes, we can better assess which areas are in need of targeted sampling in the future. Specifically, and in contrast to previous work (Girardello *et al*. 2019), we found a severe lack of sampling in the most northern regions of North America. This urgency to sample the north is further supported by the stark reality that these regions are also experiencing the most drastic impacts of climate change (Manabe and Stouffer 1980, Gauthier *et al*. 2015). Overall, several key regions should be prioritized for sampling including: (a) tundra and boreal forest, (b) tropical forests, and (c) deserts. Given the relatively low human population densities of these regions, funding directed towards establishing community science initiatives, and partnerships among organizations with interests in butterfly monitoring, will likely be critical alongside complementing these initiatives with specimen collection and focal digitization of records in these regions.

## Conclusions

Butterfly inventory completeness is not uniform across North America. Our research has revealed continuing under-sampling in regions that are facing threats from climate change as well as within specific biomes across the continent. Additionally, family level differences in sampling completeness may be driven by species traits and abundance, leading to disparities in completeness across taxa. In order to mitigate some of these issues, attention should be drawn towards establishing community partnerships of both opportunistic and structured survey systems in under-sampled regions. It is clear that community science provides a strong mechanism for alleviating sampling shortfalls and has potential to provide finer-grained views of butterfly communities, but only if such initiatives are also directed farther from regions with the densest human populations and travel infrastructure. Furthermore, additional curation and digitization of museum specimens will be critical in developing a historical backbone for analyses across time and space. Millions of specimens still remain undigitized in arthropod natural history collections (Cobb *et al*. 2019), and the continuation of funding for museum staff and biodiversity informatics infrastructure will be critical in mobilizing these data needed for ecological research, especially potential for some kinds of temporal trend analyses (Soroye et al. 2020). Supporting digitization in tandem with concerted efforts to direct community science initiatives towards under-sampled regions will move us towards unlocking the full potential of these opportunistic data in an era of global change.

## Acknowledgements

We would like to thank the countless museum staff and community scientists for their tremendous work in digitizing and documenting butterfly records from across the continent. We appreciate Michelle Duong and the Map of Life informatics team for help with Mexican distribution data. Map of Life provides visualizations of range products utilized here.

