## Supplemental Figure 1 for "Closing Gaps But Increasing Bias In North American Butterfly Inventory Completeness"

### Supplementary Material

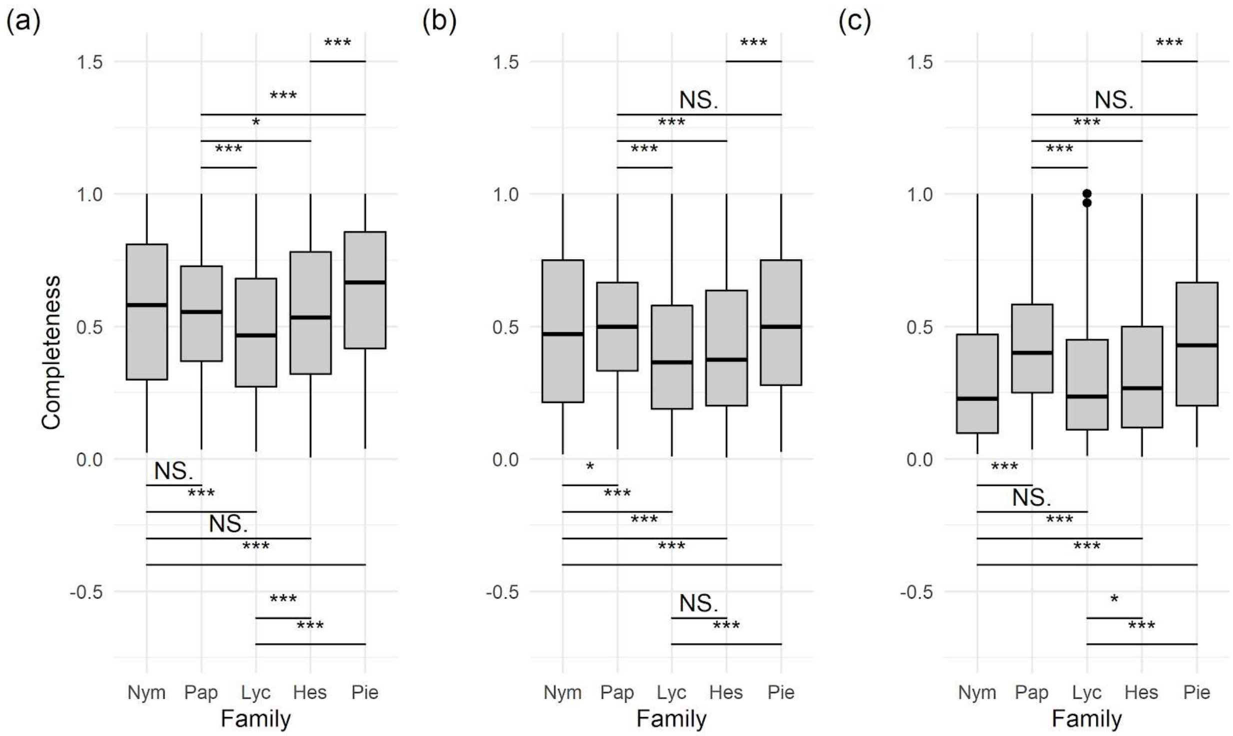


Supplemental Figure 1: Results of ANOVA post-hoc tests for average completeness in (a) the composite dataset, (b) the community observation dataset, and (c) the museum specimen dataset.

iDigBio recordsets included in this analysis are listed following the official query file: http://www.idigbio.org/portal (2020),

Query: {"filtered": {"filter": {"and": [{"terms": {"country": ["united states", "mexico", "canada"], "execution": "or"}}, {"geo_bounding_box": {"geopoint": {"bottom_right": {"lat": 10.487811882056695, "lon": -42.890625}, "top_left": {"lat": 83.7539108491127, "lon": -176.484375}}}}, {"terms": {"execution": "or", "family": ["nymphalidae", "pieridae", "papilionidae", "hesperiidae", "lycaenidae", "riodinidae"]}}]}}},

498682 records, accessed on 2020-04-02T10:36:56.211457,

contributed by 92 Recordsets, Recordset identifiers:

http://www.idigbio.org/portal/recordsets/6b5e29d3-b462-44d8-ba38-d68af5088067 (85397 records)

http://www.idigbio.org/portal/recordsets/ffafa16e-c903-4a59-8edf-b368bb6eccb3 (72909 records)

http://www.idigbio.org/portal/recordsets/d8f04cbf-f08f-4ec4-a6ae-3d473244fb16 (34017 records)

http://www.idigbio.org/portal/recordsets/92dd8c8e-c048-4f0a-9b5d-2ee627d2f553 (32644 records)

http://www.idigbio.org/portal/recordsets/1ae056ac-8834-43a8-ad44-d1148b6e075f (31682 records)

http://www.idigbio.org/portal/recordsets/6135420b-aac1-476e-bda5-e07ba8458662 (28480 records)

http://www.idigbio.org/portal/recordsets/271a9ce9-c6d3-4b63-a722-cb0adc48863f (20519 records) http://www.idigbio.org/portal/recordsets/fc628e53-5fdf-4436-9782-bf637d812b48 (20253 records)

http://www.idigbio.org/portal/recordsets/6539877e-82dc-485c-ad3d-038f383d5431 (19860 records)

http://www.idigbio.org/portal/recordsets/b985f284-eac5-4efe-8a7c-c726cdf7cf33 (18468 records)

http://www.idigbio.org/portal/recordsets/a68df423-aae9-4f4b-8a42-a36124627a53 (10083 records)

http://www.idigbio.org/portal/recordsets/55b17f44-8c6b-4dc5-a31d-d9955b425790 (9881 records)

http://www.idigbio.org/portal/recordsets/a8d88237-2f62-4ad5-b1f7-ab13ace304df (9798 records)

http://www.idigbio.org/portal/recordsets/1f9209f0-ecf2-4c3e-8ec5-bfacec1e5b9c (9472 records)

http://www.idigbio.org/portal/recordsets/b6d6cab1-f73d-4e26-86a5-e38cbe218f59 (9089 records)

http://www.idigbio.org/portal/recordsets/87017793-00dc-4f5d-b95b-09e7d17327cc (9072 records)

http://www.idigbio.org/portal/recordsets/a2aa4105-a10a-499f-8002-c6a8cb600a74 (8790 records)

http://www.idigbio.org/portal/recordsets/879d475f-4b76-4d18-8cf6-a7e5a6d44926 (6870 records)

http://www.idigbio.org/portal/recordsets/1320a827-d62a-45b1-9d26-c0aa4790a422 (6114 records)

http://www.idigbio.org/portal/recordsets/e73fedf0-90ee-4c6d-88dd-49399878fc54 (5941 records) http://www.idigbio.org/portal/recordsets/662088ed-7dc8-4043-8046-8e1035ae742f (5892 records)

http://www.idigbio.org/portal/recordsets/613232f2-7ef5-4267-b4b4-96de1c94a7e5 (5278 records)

http://www.idigbio.org/portal/recordsets/ab4b6a2b-a90a-44ce-95a1-2c44c911fcc6 (4229 records)

http://www.idigbio.org/portal/recordsets/ecb0a329-c019-4a59-abe3-fa7e72055902 (3638 records)

http://www.idigbio.org/portal/recordsets/dde625e8-cc2a-4877-9ec4-0b8a20dfded9 (3381 records)

http://www.idigbio.org/portal/recordsets/eaa5f19e-ff6f-4d09-8b55-4a6810e77a6c (3041 records)

http://www.idigbio.org/portal/recordsets/508ab646-d530-49aa-ac3c-0e2aba9a8011 (2491 records)

http://www.idigbio.org/portal/recordsets/ad5c4ec7-ed56-4d3e-881a-963af217d334 (1692 records)

http://www.idigbio.org/portal/recordsets/a6eee223-cf3b-4079-8bb2-b77dad8cae9d (1608 records)

http://www.idigbio.org/portal/recordsets/03d3b1ea-c664-4699-ac3b-420735bef8e4 (1402 records)

http://www.idigbio.org/portal/recordsets/32d433aa-9e2b-4ff9-bc55-5c3e30112207 (1293 records)

http://www.idigbio.org/portal/recordsets/34cad268-8226-4280-b637-dde38c82a29e (1223 records) http://www.idigbio.org/portal/recordsets/4f436daa-01d5-4be6-b5c3-fdd255677536 (1125 records)

http://www.idigbio.org/portal/recordsets/82541f90-fe8e-4d66-84d8-4fe515dc5533 (1094 records)

http://www.idigbio.org/portal/recordsets/2e6b6643-ebc7-4a80-a7ea-f4dd7b9c42e7 (1093 records)

http://www.idigbio.org/portal/recordsets/84006c59-fead-4b84-b3b5-cedf28f67ea9 (1040 records)

http://www.idigbio.org/portal/recordsets/ff111763-e72d-4f24-8914-b5b2dd94908c (1019 records)

http://www.idigbio.org/portal/recordsets/3c919328-94fd-4657-b81d-21f4707253ed (884 records)

http://www.idigbio.org/portal/recordsets/2c662e9e-cdc6-4bbf-93a5-1566ceca1af3 (820 records)

http://www.idigbio.org/portal/recordsets/cb4882a8-19d3-41a1-9373-c1a868b5ac9b (791 records)

http://www.idigbio.org/portal/recordsets/9fab87bf-c5a7-4296-acdc-f22859cfe620 (657 records)

http://www.idigbio.org/portal/recordsets/6075f4b7-4242-4886-be7d-391614fffe41 (618 records)

http://www.idigbio.org/portal/recordsets/b80d24e8-17ff-4092-88d7-7e5ad11c117a (564 records)

http://www.idigbio.org/portal/recordsets/8eabbc48-2b30-419d-bd8f-eece9185eca1 (529 records)

http://www.idigbio.org/portal/recordsets/65007e62-740c-4302-ba20-260fe68da291 (499 records)

http://www.idigbio.org/portal/recordsets/7fcdca8e-7469-480c-8516-cce4e24c37c9 (462 records)

http://www.idigbio.org/portal/recordsets/f1512610-8631-475c-875a-a634191a9715 (390 records)

http://www.idigbio.org/portal/recordsets/f93c403b-d0f0-4be1-8a4b-ed9aa3b513e3 (351 records)

http://www.idigbio.org/portal/recordsets/69037495-438d-4dba-bf0f-4878073766f1 (302 records)

http://www.idigbio.org/portal/recordsets/dce98228-c00d-4532-a614-c20384e34e24 (295 records)

http://www.idigbio.org/portal/recordsets/a47eacbb-22d0-4a8b-8853-989b26fd8290 (216 records) http://www.idigbio.org/portal/recordsets/bf897c17-06cc-48c1-a7cc-f41b45166880 (215 records)

http://www.idigbio.org/portal/recordsets/9354db6c-4019-4351-a822-cb87b1b73a44 (213 records)

http://www.idigbio.org/portal/recordsets/cf42855e-a54a-4488-a79e-beac086ba1d4 (157 records)

http://www.idigbio.org/portal/recordsets/a5ec0be5-1c6f-448a-b75d-76085fe3ff20 (143 records)

http://www.idigbio.org/portal/recordsets/53feaa83-e3b6-4ad3-8597-293b153e7548 (115 records)

http://www.idigbio.org/portal/recordsets/da67ebd9-52de-444d-b114-e23c03111ac6 (103 records)

http://www.idigbio.org/portal/recordsets/568e209f-d072-4fd6-8b64-27954b0fd731 (89 records)

http://www.idigbio.org/portal/recordsets/5aef068f-efd3-4851-a623-f542e97350cd (73 records)

http://www.idigbio.org/portal/recordsets/58d0090c-167e-4d7a-8ad5-c50681626692 (65 records)

http://www.idigbio.org/portal/recordsets/f885076c-a7ae-4a7b-9769-3f733c6a9ecf (47 records)

http://www.idigbio.org/portal/recordsets/cb65cf5e-07b8-4d53-a91a-7dce9b8ccf80 (40 records)

http://www.idigbio.org/portal/recordsets/0395f3c4-0277-43b6-a36d-e07720588790 (24 records)

http://www.idigbio.org/portal/recordsets/7f7939c8-5276-4fbb-b12c-de89a5a8044f (19 records)

http://www.idigbio.org/portal/recordsets/2292f5d5-d39f-4944-be72-fa5dd62f581c (15 records)

http://www.idigbio.org/portal/recordsets/6bb853ab-e8ea-43b1-bd83-47318fc4c345 (15 records)

http://www.idigbio.org/portal/recordsets/3617a6a3-d384-48de-948d-2d1e1c54e090 (14 records)

http://www.idigbio.org/portal/recordsets/e597a4f6-2be7-4538-8007-dec480dc4d6f (10 records)

http://www.idigbio.org/portal/recordsets/95ecb448-3c1f-4145-8565-4f6d51beb62c (9 records)

http://www.idigbio.org/portal/recordsets/59422682-15ba-47e1-99e2-1ef69f7bdd9a (8 records)

http://www.idigbio.org/portal/recordsets/db4bb0df-8539-4617-ab5f-eb118aa3126b (7 records)

http://www.idigbio.org/portal/recordsets/656d1c9a-cbb7-4dde-ac24-62323af5b831 (6 records)

http://www.idigbio.org/portal/recordsets/56879e73-bf9d-4bd9-a7a4-4f2f940d0f62 (6 records) http://www.idigbio.org/portal/recordsets/cfe5283d-89df-4662-8c45-8f0c76e90107 (6 records)

http://www.idigbio.org/portal/recordsets/5939498b-0fcc-42ca-929b-99f4b2c5e1aa (4 records)

http://www.idigbio.org/portal/recordsets/d315c4a3-0bee-49d1-8d03-726358937cde (4 records)

http://www.idigbio.org/portal/recordsets/9d9b81f1-3a9a-4515-b741-a145293f1fef (3 records)

http://www.idigbio.org/portal/recordsets/5e893602-84ca-4c8c-bac1-99111c777582 (2 records)

http://www.idigbio.org/portal/recordsets/d621e959-2633-4ec1-a2a2-5d97cd818b47 (1 records)

http://www.idigbio.org/portal/recordsets/10eea9b5-e08f-4c07-adf6-561aea89ae09 (1 records)

http://www.idigbio.org/portal/recordsets/f778ecc0-8371-49d5-9ab1-9d75f0b76fad (1 records)

http://www.idigbio.org/portal/recordsets/2b89de41-42bd-46c6-ab61-d386f855f7fb (1 records)

http://www.idigbio.org/portal/recordsets/347579f4-d44a-4c8e-a578-09c2a8132573 (1 records)

http://www.idigbio.org/portal/recordsets/e3063807-1c2e-4eed-b5b9-170e26e8b565 (1 records)

http://www.idigbio.org/portal/recordsets/25cd5e12-7830-4f46-bf6d-9b6deb706f44 (1 records)

http://www.idigbio.org/portal/recordsets/4146072c-ed77-4bed-a03a-8363982a91e8 (1 records)

http://www.idigbio.org/portal/recordsets/513dd822-f211-42c3-9af2-ee3c798cd1b8 (1 records)

http://www.idigbio.org/portal/recordsets/7df4be6b-67e9-4c05-8cd9-f546bdcc54d0 (1 records)

http://www.idigbio.org/portal/recordsets/b5f4526b-f4fb-4d90-8ce0-975e0cda8ff6 (1 records)

http://www.idigbio.org/portal/recordsets/137ed4cd-5172-45a5-acdb-8e1de9a64e32 (1 records)

http://www.idigbio.org/portal/recordsets/5889291d-9105-4740-a30f-2d9d2469c264 (1 records)

http://www.idigbio.org/portal/recordsets/837d99c6-3045-4ba3-8951-643ddb3d6676 (1 records)
